# RNA polymerase I passage through nucleosomes depends on its lobe binding subunits

**DOI:** 10.1101/823658

**Authors:** Philipp E. Merkl, Michael Pilsl, Tobias Fremter, Katrin Schwank, Christoph Engel, Gernot Längst, Philipp Milkereit, Joachim Griesenbeck, Herbert Tschochner

## Abstract

RNA polymerase I (Pol I) is a highly efficient enzyme specialized to synthesize most of the ribosomal RNA. After nucleosome deposition at each round of replication the Pol I transcription machinery has to deal with nucleosomal barriers. It was suggested that Pol I-associated factors facilitate chromatin transcription, but it is not known whether Pol I has an intrinsic capacity to transcribe through nucleosomes. Here we used *in vitro* transcription assays to study purified Pol I of the yeast *S. cerevisiae* and Pol I mutants in comparison to Pol II and Pol III to pass a nucleosome. Under identical conditions, purified Pol I and Pol III, but not Pol II, were able to transcribe nucleosomal templates. Pol I mutants lacking either the heterodimeric subunit Rpa34.5/Rpa49 or the C-terminal part of the specific subunit Rpa12.2 showed a lower processivity on naked DNA templates, which was even more reduced in the presence of a nucleosome. The contribution of Pol I specific subunit domains to efficient passage through nucleosomes in context with transcription rate and processivity is discussed.

## Introduction

Eukaryotic RNA polymerases (RNAPs) share many structural and functional features, but are also are highly specialized transcription machineries. The molecular nature of their specialisation is presently subject of intensive investigations (see as reviews [1–3]. In most eukaryotes, RNA polymerase I (Pol I) transcribes only one gene locus, the multi copy ribosomal (r)RNA genes. Even in exponentionally growing cells only a fraction of the more than 100 copies of rRNA genes are transcribed [4]. More than 100 transcribing Pol I molecules can be packed on one rRNA gene [5,6]. The dense packing of elongating Pol I molecules is probably not compatible with a regular nucleosomal structure. In agreement with this, most nucleosomes are depleted from transcribed rRNA genes, resulting in an open chromatin structure at transcribed rRNA genes [7,8]. The balance between the transcriptionally active (open) and nucleosomal (closed) rRNA chromatin states is due to a dynamic equilibrium of transcription-dependent removal and replication dependent assembly of nucleosomes [9]. Although it was shown that Pol I transcription is needed to establish the open chromatin structure, it is still unclear how nucleosomal barriers affect Pol I transcription and if components of the basal Pol I machinery support chromatin transcription. Along this line, it is possible that Pol I subunits influence transcription of nucleosomal templates.

Whereas knowledge about Pol I transcription of nucleosomal templates is rather sparse, transcription of nucleosomal templates by purified RNA polymerase II was studied in molecular detail (see as reviews[10–12]). Several factors were described to act on nucleosomes to modulate the barrier and facilitate Pol II passage (see as review [10]). One of them, FACT (facilitates chromatin transcription), was also suggested to support Pol I transcription through nucleosomes [13]. Other Pol II associated factors are able to reduce intrinsic stalling of Pol II on chromatin templates thereby reducing the amount and rate of backtracked and arrested enzymes (see as review[11]). Among these factors are the Pol II transcription factors TFIIF and TFIIS which are structurally and functionally related to the heterodimeric Pol I subunit Rpa34.5/Rpa49 and subunit Rpa12.2, respectively [14][15,16]. Interplay between the Pol II factors TFIIF and the RNA cleavage factor TFIIS was reported to promote transcript elongation and increase transcription through nucleosomes in a synergistic manner [17,18]. Whether the Pol I counterparts play a role in nucleosomal transcription remains to be elucidated.

The heterodimer Rpa34.5/Rpa49 of Pol I corresponds to Rpc37/Rpc53 of Pol III [3] and associates together with the N-terminal part of Rpa12.2 to the Pol I lobe structure, thereby forming a lobe binding module (hereafter called lb-module). The heterodimer Rpa34.5/Rpa49 consists of three subdomains: a dimerization module formed by Rpa34.5 and the N-terminal part of Rpa49 (full length Rpa34.5 and aa 1-110 of Rpa49), the A49 linker (aa 105-187 of Rpa49), and the C-terminal part of Rpa49 (aa 187-415, hereafter called Rpa49CT). The dimerization module binds to the ‘lobe’ and ‘external’ domains of the second largest Pol I subunit Rpa135 on the core module side [15,19,20] and occupies a similar position as the TFIIF dimerization module (yeast Tfg1/Tfg2) in association with Pol II [21,22]. The C-terminal part of Rpa49 forms a tandem winged helix (tWH) similar to the small TFIIE subunit Tfa2 [16,23] or in RPC37 [24]. This domain is flexibly attached to the Pol I core in structures of dimeric and free Pol I or in elongation complexes [19,20,25,26]. Similar to TFIIF the Pol I subcomplex Rpa34.5/Rpa49 was suggested to enhance RNA elongation [14,16,27,28] and to stabilize the initiation complex [15,29], presumably contributing to a high rate of Pol I loading [30]. The Pol I lb-module contains also the N-terminal part of subunit Rpa12.2 (amino acids (aa) 1 to 85, hereafter called Rpa12ΔC)). The Rpa12.2 N-terminus is homologous to the N-terminal domains of Pol II subunit Rpb9 and Pol III subunit Rpc11[2]. In contrast, the C-terminal part of Rpa12.2 (aa 85 to 125, Rpa12CT) is homologous to the Pol II RNA cleavage factor TFIIS and to the C-terminus of Pol III subunit Rpc11 [14,31]. As TFIIS in the Pol II system, Rpa12.2CT reaches through a pore to the active site of Pol I and supports the intrinsic RNA cleavage activity of the polymerase [19,20,32]. Deletion of Rpa12.2 results in growth inhibition at elevated temperature, sensitivity to nucleotide-reducing drugs, inefficient transcription termination, transcription errors and hampers the assembly of the Pol I enzyme [33–36]. In summary, Rpa12.2 as well as Rpa34.5/Rpa49 promote the elongating Pol I. The lobe associated N-terminal domain of Rpa12.2 is important for proper Pol I assembly, whereas the C-terminal domain supports RNA cleavage.

To get insights whether and how the lobe associated subunits influence transcription of nucleosomal templates we analysed the function of the Rpa34.5/Rpa49 heterodimer and Rpa12.2 using defined promoter-dependent and -independent *in vitro* transcription systems. We studied effects of suboptimal transcription conditions for Pol I to pass a nucleosome and compared purified Pol I with Pol II and Pol III in nucleosomal transcription. Our investigations emphasize the roles of Rpa49LCT (containing the linker and the tWH domain of Rpa49, i.e. aa 105-415) and Rpa12CT for assuring Pol I transcription through an *in vitro* assembled nucleosome.

## Materials and Methods

### Yeast strains, plasmids, oligonucleotides and construction of transcription templates

Oligonucleotides, plasmids and yeast strains used in this work are listed in Appendix Tables S1, S2 and S3. Molecular biological methods and transformation of yeast cells were performed according to standard protocols [37][38,39]. Transcription templates were generated by PCR using the indicated oligonucleotides and plasmids. Plasmid sequences are available upon request. To purify Pol I from yeast, cells were grown at 30°C in either YPD (2% w/v) peptone, 1% (w/v) yeast extract, 2% glucose) or on YPG (2% w/v) peptone, 1% (w/v) yeast extract, 2% galactose).

### Purification of proteins

Wild-type RNA polymerases I, II and III (Pol I, II and III) were purified from yeast strains y2423 (Pol I), y2424 (Pol II), y2425 Pol III, (Appendix Table S1) via the protein A affinity tag according to [40]. Purification of mutant Pol I y2670 (ΔRpa49), y2679 (ΔRpa12) and y2679 transformed with plasmid 1839 (Rpa12ΔCT) was according to [41]. Recombinant Rpa34/Rpa49, the dimerization domain Rpa34.5/Rpa49NTL (Rpa34/(Rpa49 aa 1-186) and Rpa49LCT (aa 112-415) were purified from *E.coli* according to the protocol of [41].

### Transcription elongation assays

Promoter-dependent elongation assay. Pulse-labelling of G-less transcripts on promoter-containing templates and subsequent analysis were performed according to [41,42] with some modifications. Templates were generated by PCR amplification of plasmid 2148 or 2313 using the forward primer 4228 and the reverse biotinylated primers 4018 or 4220. The resulting DNA fragments could be transcribed into 1 kb and 2kb long RNAs, respectively. If elongation should be measured during 5 time points including the starting point at time point 0, the following assay was performed. 1.5 ml reaction tubes (Sarstedt safety seal) were placed on ice. 5 × 0.5 to 1 μl template (50 – to 100 ng DNA) (templates 2148 or 2313) immobilized to magnetic beads was added, which corresponds to a final concentration of 5 – 10 nM per transcription reaction (25 μl reaction volume). 5 - 10 μl CF (0.5 to 1 pmol/μl; final concentration 20 - 40 nM) 2 μl Rrn3 (final concentration 70 nM) and 5 - 15 μl Pol I (final concentration 4 - 12 nM) were added to the tube. Where indicated, domains of the Rpa34.5/Rpa49 heterodimer were added before transcription was started (final concentration 50 nM). 20 mM HEPES/KOH pH 7.8 were added to a final volume of 125 μl. Transcription was started adding 5 × 12.5 μl transcription buffer 2 x without GTP (40 mM HEPES/KOH pH 7.8, 20 mM MgCl_2_, 10 mM EGTA, 5 mM DTT, 300 mM potassium acetate, 0.4 mM ATP, 0.4 mM UTP, 0.02 mM CTP, 0.3 μCi ^32^P-CTP). The samples were incubated at 24°C between 5 and 30 min at 400 rpm in a thermomixer. After placement on ice, the supernatant was separated from streptavidin-covered magnetic beads and 1-3 washing steps using wash buffer W1 (20 mM HEPES/KOH pH 7.8, 10 mM MgCl_2_, 5 mM EGTA, 2.5 mM DTT, 150 mM potassium acetate, 0.02 mM ATP, 0.02 mM UTP, 0.02 mM CTP followed). The magnetic beads were resuspended in 150 μl wash buffer W1. 25 μl were removed and added to 200 μl Proteinase K buffer (0.5 mg/ml Proteinase K in 0.3 M NaCl, 10 mM Tris/HCl pH 7.5, 5 mM EDTA, 0.6% SDS) (time point 0). To the remaining 125 μl either 12.5 μl or 1.25μl 10xNTPs (ATP, GTP, UTP, CTP concentration each 2 mM) and Heparin (20 ng/μl) was added and immediately mixed. 25 or 27 μl (in case of adding 12.5 ul NTPs) aliquots were removed at the time points indicated and added to 200 μl Proteinase K buffer. The samples were incubated at 30°C for 15 min at 400 rpm in a thermomixer. 700 μl Ethanol p.a. were added and mixed. Nucleic acids were precipitated at −20°C over night or for 30 min at −80°C. The samples were centrifuged for 10 min at 12.000g and the supernatant was removed. The precipitate was washed with 0.15 ml 70% ethanol. After centrifugation, the supernatant was removed and the pellets were dried at 95°C for 2 minutes. RNA in the pellet was dissolved in 12 μl 80% formamide, 0.1 TBE, 0,02% bromophenol blue and 0.02% xylene cyanol. Samples were heated for 2 min under vigorous shaking at 95°C and briefly centrifuged. After loading on a 6% polyacrylamide gel containing 7 M urea and 1 x TBE RNAs were separated applying 25 watts for 30 - 40 min. After 10 min rinsing in water the gel was dried for 30 min at 80° C using a vacuum dryer. Radiolabelled transcripts are visualised using a PhosphoImager. For quantification, signal intensities were calculated using Multi Gauge (Fuji).

Tailed template elongation assay. The transcription were performed as described above on PCR-derived templates containing a 3’overhang. Biotinylated oligos downstream of the tail in reverse direction (oligos 4018 and 4220 for 1 kb and 2 kb transcripts, respectively) and an oligo containing a Nb.BsmI (NEB) nicking site (oligo 4220a) were used to generate templates of 1 and 2 kb using plasmid 2316 as template. PCR products were cut with Nb.BsmI and heat inactivated at 80°C for 20 min. After 10 min, a competitor oligo 4220a with the same sequence as the 24 nt 3’overhang was added in excess to anneal with released cleaved 5’Nb.BsmI fragment. The DNA was ethanol precipitated and resuspended in water. Tailed templates allowed non-specific initiation of all RNA polymerases without the addition of initiation factors.

### Transcription of nucleosomal templates

Preparation of nucleosomal templates: Plasmid 1253 containing one 601 nucleosome binding site and plasmid 1573 without 601 binding site were linearized with EcoRV and PvuII followed by incubation for 20 min at 65°C to inactivate EcoRV. The DNA was ethanol precipitated, washed with 70% EtOH and resuspended in RNase free water. The the DNA was digested with Nb.BsmI at 65°C to obtain a 3’overhang. PvuII and Nb.BsmI were heat inactivated at 80°C for 20 min. After 10 min, a competitor oligo 2207 with the same sequence as the 24 nt 3’overhang was added in excess to anneal with released cleaved 5’Nb.BsmI fragment. The DNA was ethanol precipitated and resuspended in water. The length of the resulting reference transcript was 240 nt, the length of the 601 sequence was 468 nt. For promoter-dependent transcription, one template was generated by PCR using plasmids 1247 (with 601 sequence) and oligos 4228 and 4233, which was transcribed into a 191 nt long transcript. The reference template was generated by PCR using plasmid 1959 and oligos 4228 and 4234. This template was transcribed into a 468 nt long RNA.

Chromatin was reconstituted on templates for *in vitro* transcription containing one 601 nucleosome positioning sequences [43]. Purification and assembly of chicken histones were performed according to [44]. Dialysis chambers were prepared from siliconized microreaction tubes (Eppendorf) by clipping off the conical part and perforation of the cap with a red-hot metal rod (ø 0.5 cm). The dialysis membrane (molecular weight cutoff 6.8kDa) was pre-wet in high salt buffer and fixed between tube and cap. The dialysis chambers were put in a floater in a bucket containing 300 ml high salt buffer, air bubbles were removed and the mixture of histones and DNA was applied to the chambers. Dialysis from 2 M NaCl to 0.23 M NaCl was performed over night at RT with constant stirring and at a low salt buffer flow rate of 200 ml/h (3 l total). Upon completion, the assembly solution was transferred to a siliconized tube, its volume was measured and chromatin was stored at 4°C. To determine the assembly success, 5 μl of the reaction were supplemented with 1μl loading buffer and analyzed on a 6% polyacrylamide, 0.4xTBE gel (Appendix Fig. S1). Gels were stained in 0.4x TBE containing ethidium bromide for 15min, washed in 0.4x TBE for 15 min and analysed on a FLA3000 imager (Fuji).

To determine optimal assembly conditions, histones were titrated to the DNA in the following ratios 0,4:1; 0.6:1; 0.8:1 and 1:1 (DNA/histones). 10 μg of BSA and 5 μg template were added. The concentration of the chicken histones was 0.8 mg/ml. Between 2 and 5 μg chicken histones were assembled in a total volume of 50 μl high salt buffer (10 mM Tris, pH 7.6, 2 M NaCl, 1 mM EDTA, 0.05% NP-40, 2 mM β-mercaptoethanol). In most reaction series the ratios 0.6:1 or 0.8:1 were used. For the tailed template analysis depicted in Appendix Fig. S4A/B the nucleosome containing template was generated by PCR using plasmid (2316) and oligo 2115 and 4233. The reference template was generated by PCR using plasmid 1573 and oligos 2115 and 4234. 3’overhangs were produced as described above using cleavage by Nb.BsmI. Each transcription reaction contained 50-100 ng of both nucleosome-containing and naked template.

Promoter-dependent transcription reactions were performed as described [41]. Each reaction contained 50-100 ng of a nucleosome-containing (1247) and 50 - 100 ng of naked template (1959). Templates of appropriate sizes were generated by PCR reactions of different sizes using oligos 4228, 4233, 4228 and 4234.

If promoter-dependent nucleosomal transcription elongation was performed with either reduced (20 μM) NTP concentration or at 0°C the transcription assay was modified. Promoter-dependent transcription was performed on templates with and without attached nucleosomes using 70 nM Rrn3, 20 nM CF and 4 nM Pol I WT. The templates contained a 22nt long stretch without guanines after the start site on the coding strand. To achieve sufficient and comparable elongation competent Pol I complexes, transcription was started using 200 μM ATP, UTP and 10 μM α^32^P-CTP, but no GTP which should result in stalled ternary complexes 22 nt downstream of the Pol I start site. After 5 min the transcription reaction was either 10 x diluted with transcription buffer containing 200 μM ATP, UTP and 10 μM ^32^P labeled CTP and 200 μM GTP or transcription buffer containing 20 μM GTP and 10 μM α^32^P-CTP but no ATP and UTP which should result in a final NTP concentration of 20 μM. The same initiation procedure (200 mM ATP, UTP, α^32^P-CTP, but no GTP, at 24 °C) was used to study elongation at different temperature. After 5 min the transcription reaction was 10 x diluted with either 24°C or 0°C cold transcription buffer containing 200 μM ATP, UTP and 10 μM ^32^P labeled CTP and 200 μM GTP. Transcription elongation was stopped after 10 min.

To analyse template protection by nucleosomes before and after transcription (see Appendix Fig. S5) templates containing 601 sequence were generated by PCR using oligos 1228 and the biotinylated oligo 4232 on template 1247. Nucleosomes were assembled as described above. After assembly templates were immobilized on streptavidin covered magnetic beads. Each 0.6 μg assembled DNA (25 μl) and not assembled DNA were incubated with 10 μl streptavidin beads (Dynabeads, 10 mg/ml) washed twice with 0.4 M NaCl, 0.15 mM EDTA, 5 mM Tris/HCl pH 7.5, 0.02 mg/ml BSA and once with 20 mM HEPES pH 7.8 and resuspended in 20 μl TE. 1.3 μl of the templates were used for promoter-dependent and promoter-independent transcription resulting in a 242 nt long transcript.

### Quantification of transcription of nucleosomal templates

For each reaction signal intensities of the bands corresponding to the two different transcripts in one lane were determined using a FLA3000 imager (Fuji). Signal intensities of transcripts derived from 601 templates with and without assembled nucleosome were divided by the signal intensities of transcripts derived from the reference template. This yielded the normalized transcript levels derived from nucleosome-free 601 and reference template. Then the quotient of full length transcript derived from the nucleosomal 601 template versus reference transcript was calculated (normalized level of nucleosomal transcripts). Finally, the ratio of the normalized 601 full length transcript signal intensities in presence and absence of the nucleosome was calculated, which was taken as measure for the efficiency of transcription through a nucleosome. The signal of the transcript from template 601 without nucleosome to the reference transcript was set to 1.

### Restriction enzyme digestion of transcription templates and quantification of nucleosomal protection by quantitative PCR

Transcription templates coupled to magnetic beads in a total volume of 25µl transcription buffer were supplemented with 175µl of 1x CutSmart buffer (NEB) containing 25u of BsiWI-HF (NEB). The bead suspension was incubated for 1h at 37°C in a thermomixer under shaking with 800 rpm. The reaction was terminated by the addition of 200µl of IRN buffer [50mM Tris–HCl (pH 8), 20mM EDTA, 500mM NaCl]. Proteinase K and SDS were added to a final concentration of 0.33 mg/ml and 0.5%, respectively. The incubation was continued for 1 h at 56°C. After phenol/chloroform extraction, DNA was precipitated with ethanol in the presence of 40 µg of glycogen as a carrier. The DNA was suspended in 50 µl of TE buffer [10mM Tris–HCl (pH 8), 1mM EDTA]. The DNA was analyzed by quantitative PCR using SYBR green I dye (Roche) for DNA detection with a Rotor-Gene 6000 system (Corbett Life Science/Qiagen). Primer pairs used for amplification were 2976, 3117, 4023, 4231, and 4273 (Table 3). All samples were analysed in triplicate PCR reactions. Relative DNA amounts in each sample were determined using the comparative analysis software module of the Rotor-Gene 6000 series software

## Results

### Pol I and III can transcribe a nucleosomal template

In contrast to Pol I and Pol III, the lb-module, is absent in Pol II. The lb-module of Pol I and Pol III is formed by subunits Rpa34.5/Rpa49, Rpa12.2 and Rpc37/Rpc53, Rpc11 respectively. It was previously published that purified Pol III can transcribe through a nucleosome [45] whereas Pol II is inhibited without inclusion of additional factors [46,47]. Stimulation of Pol II transcription through nucleosomes was achieved in the presence of several elongation factors including TFIIS, FACT [47], Spt4/5 [48] and Paf1C [49]. Furthermore, a synergistic positive effect of TFIIS and TFIIF on nucleosomal transcription by Pol II was reported [18]. Interestingly, FACT supported as well Pol I transcription through nucleosomes [13]. Here, the three purified nuclear RNA polymerases from yeast were directly compared in their ability to transcribe nucleosomal templates. Pol I, Pol II and Pol III were purified using the same one-step affinity purification procedure and identical buffers [40]. The purified enzymes were free of detectable cross-contamination of transcription factors or RNA synthesizing enzymes [40]. Nucleosomes were assembled by salt dialysis at the 601 nucleosome positioning sequence [43] on a tailed template carrying a 3’-ssDNA overhang (see Appendix Fig. S1 and Material and Methods). The use of tailed templates allowed promoter independent transcription initiation by all three polymerases [40]. For thorough quantification of the efficiency of RNA synthesis from a nucleosomal template, transcription reactions were performed in the presence of an additional nucleosome-free reference template, serving as an internal control. Transcription reactions were performed with 7.5 nM of the respective polymerases at identical salt concentrations (150 mM potassium acetate). All three polymerases were able to transcribe reference templates and/or the 601 sequence containing templates without nucleosomes quite efficiently (Fig. 1, lanes 2, 4, 5, 7, 9, 10, 12, 14, 15). Differences in signal intensities between the generated RNAs were due to the use of a universal buffer system that may not be ideal for all enzymes. The presence of a nucleosome on the DNA template reduced the efficiency of transcription. However, the level of inhibition differed significantly between the three polymerases. Whereas a major fraction of Pol I and Pol III could transcribe the nucleosomal 601 sequence (Fig. 1A, lanes 1, 3, 11, 13), full-length transcripts from nucleosomal templates were almost not detectable in Pol II-dependent reactions (Fig. 1A, lanes 6 and 8). Upon longer exposure times additional smaller transcripts became visible in lanes 6 (Appendix Fig. S2A). These smaller transcripts could be products of abortive or arrested transcription when Pol II encountered the nucleosome. This indicated that at least a minor fraction of Pol II initiated transcription from the nucleosome-decorated tailed template but was arrested or displaced upon collision with a nucleosome. Quantification revealed about 40% reduced RNA synthesis by Pol I and Pol III in the presence of a nucleosome and almost complete loss of Pol II-derived transcripts (Fig. 1C). These results suggest that Pol I and Pol III, but not Pol II are capable to transcribe through a nucleosomal barrier without additional factors.

**Fig 1.**
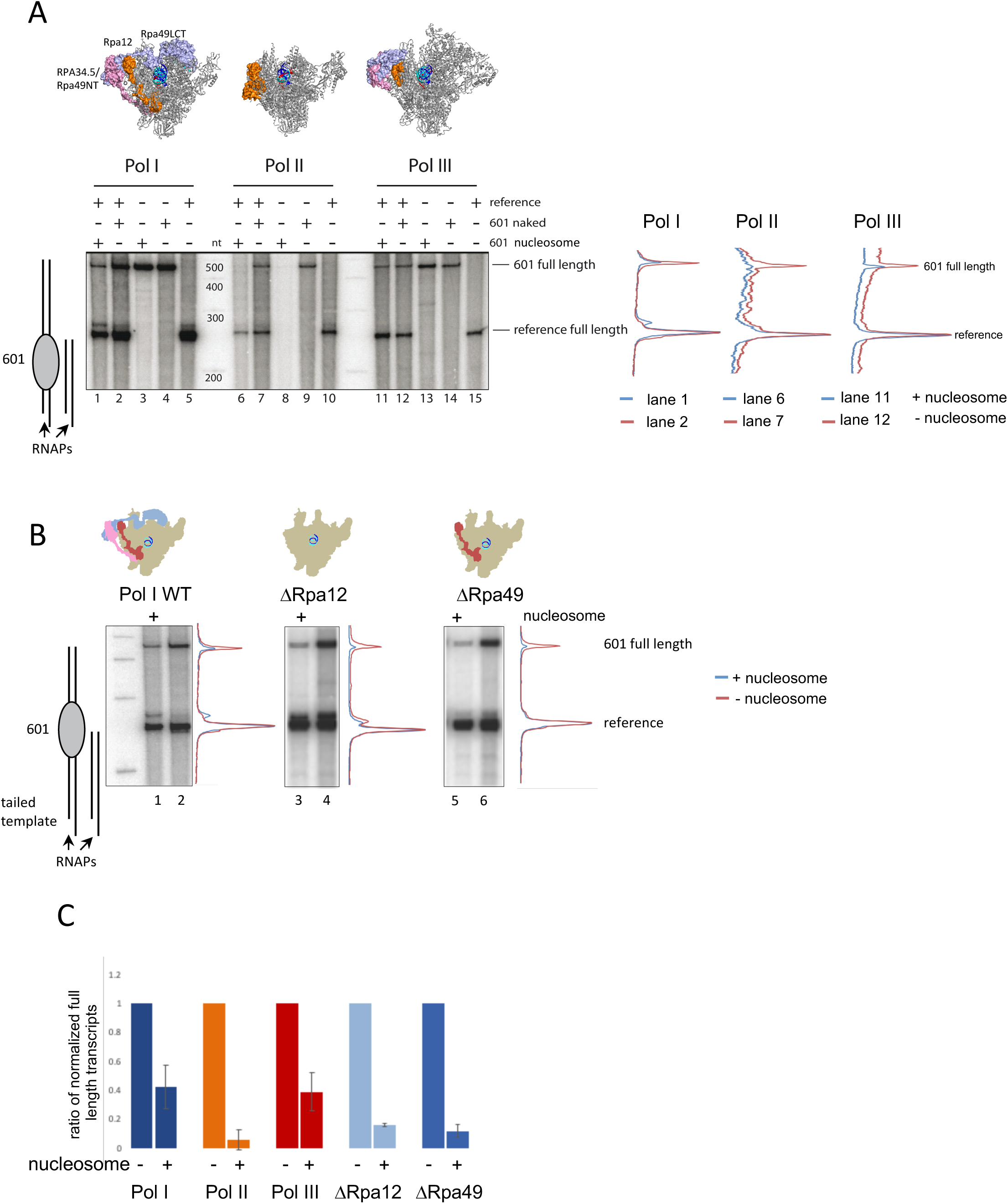
Efficiency of transcription through a nucleosome in tailed templates differs between Pol I, Pol II and Pol III and is reduced in Pol I mutants lacking lobe-binding subunits. A) Nucleosomes were assembled on tailed templates containing a 601 nucleosome positioning sequence (601) (derived from plasmid 1253) as described in material and methods (see also Appendix Fig.S1). Transcription reactions were performed in the presence of a 601-containing and a 601-free reference template. The cartoon on the left indicates the length of the transcripts derived from the 601-containing and reference template. *In vitro* transcription assays were performed as described using 7.5 nM of purified RNA polymerases, identical buffer conditions and 10 nM of template 601 with or without nucleosomes and the reference template containing neither nucleosomes nor the 601 sequence. Structures of the three nuclear yeast RNA polymerases with the lobe binding subunits Rpa34.5/Rpa49 (Pol I) and Rpc37/Rpc53 (Pol III) in magenta/light blue and Rpa12.2 (Pol I) and Rpc11 (Pol III) and Rpb9 (Pol II) in orange are modelled according to pdb 5M3F [26], pdb 4C2M [20], pdb 5F8J [24], 8W6S [58] and pdb 1Y1W [73]. Transcripts were analyzed on a 6% polyacrylamide gel containing 6 M urea and detected using a PhosphorImager. On the right side the transcript profiles of the incicated lanes are shown. The value of the reference transcript was set to 1. Length of reference and 601 transcript is 240 nt and 468 nt, respectively. B) Pol I mutants lacking either the heterodimeric subunit Rpa34.5/Rpa49 or subunit Rpa12.2 are hampered in efficient transcription through a nucleosome in tailed template assays. Transcription reactions were performed on tailed templates using 5 nM Pol I WT, 20 nM Pol I ΔRpa49 or Pol I ΔRpa12.2 as in A). Transcript profiles of each lane are shown on the right. The value of the reference transcript was set to 1. The cartoon on the left indicates the length of the transcripts derived from the nucleosomal and reference template. The cartoons on top show Pol I models with or without Rpa34.5 (magenta), Rpa49 (light blue) and Rpa12.2 (orange). C) Quantification of the experiments shown in A and B. Signal intensities of 601 full length transcripts (Fig. 1A, lanes 1/2, 6/7 and 11/12 of Pol I, Pol II and Pol III)(Fig. 1B, lanes 1/2, 3/4 and 5/6 of Pol I WT, Pol I ΔRpa12.2 and Pol I ΔRpa49) were normalized to reference full length transcript signal intensities in the respective lanes. Then, the ratio of the normalized 601 full length transcript signal intensities in presence and absence of the nucleosome was calculated. Pol I WT (n=8), Pol II (n=5), Pol III (n=5), Pol I ΔRpa49 (n=5), Pol I ΔRpa12 (n=3). Mean values were plotted including error bars of the standard deviation.

### Lobe-binding Pol I subdomains are important for efficient transcription of nucleosomal templates *in vitro*

In Pol II-dependent transcription, TFIIF together with the cleavage stimulating factor TFIIS promotes elongation [17] and RNA synthesis through nucleosomes [18]. Since Rpa34.5/Rpa49 and Rpa12.2 share structural features with TFIIF and TFIIS, one could hypothesize that these subunits might facilitate transcription through a nucleosome. Therefore, we investigated whether absence of Rpa12.2 and/or the Rpa34.5/Rpa49 heterodimer had an impact on Pol I transcription of a nucleosomal template.

To study the influence of Pol I lb-module subunits on *in vitro* transcription, wildtype and mutant Pol I were affinity-purified from yeast through protein A-tagged subunit Rpa135 using the same salt conditions [40]. Purified wild-type (WT) Pol I contained all 14 subunits, ΔRpa49 Pol I lacked the heterodimeric subunit Rpa34.5/Rpa49, ΔRpa12.2 Pol I lacked subunit Rpa12.2 and the heterodimer Rpa34.5/Rpa49, and Rpa12ΔC Pol I contained the N-terminal part of Rpa12.2 and the Rpa34.5/Rpa49 heterodimer (Appendix Fig. S3A, B), in good correlation with previous analyses [34].

Consistent with a role of the lobe binding subunits in transcription through a nucleosome, both, ΔRpa12.2 Pol I and ΔRpa49 Pol I could not as efficiently transcribe nucleosomal templates as the WT enzyme (Fig. 1B). In contrast to WT Pol I in which efficiency of nucleosomal transcription was reduced to about 40% compared to transcription without assembled nucleosome, Pol I lacking Rpa34.5/Rpa49 and/or Rpa12.2 less than 20% fully extended nucleosomal transcripts could be measured (Fig. 1C).

Rpa34.5/Rpa49, Rpa34.5/Rpa49 or individual subdomains were recombinantly expressed in *E. coli* and purified (Appendix Fig. S3C), to test if their presence could support ΔRpa49 Pol I transcription of a nucleosomal template. As a control, we verified that the purified heterodimer alone contained no significant RNA synthesizing activities (Fig. 2A, lanes 5 and 6). Addition of increasing amounts of the purified heterodimer increased transcriptional activity of ΔRpa49 Pol I on both reference and nucleosomal template (Fig. 2A), However, nucleosomal transcription was significantly more stimulated (Fig. 2A, lanes 7-14, and quantification). We next investigated the individual contributions of Rpa34.5/Rpa49 domains to support transcription through a nucleosome. Thus, either the dimerization module composed of the N-terminus of Rpa49 including the linker (Rpa49NTL, aa 1 to 186) in complex with Rpa34.5, or the C-terminal domain of Rpa49 including the linker (Rpa49LCT, aa 112 to 415) were tested in *in vitro* transcription reactions together with ΔRpa49 Pol I. Similar to Rpa34.5/Rpa49, addition of Rpa49LCT increased the transcript levels from both the reference template and the nucleosomal template. (Fig. 2B; Appendix Fig. S4A-C). Transcript levels varied in dependency of the quality of the preparation of the nucleosomal DNA matrix and the quality of the Pol I preparation. Adding the dimerization module Rpa34.5/Rpa49NTL alone to Pol I ΔRpa49 did not show significant enhancement of transcription (Fig. 2B)(Appendix Fig. S4).

**Fig. 2.**
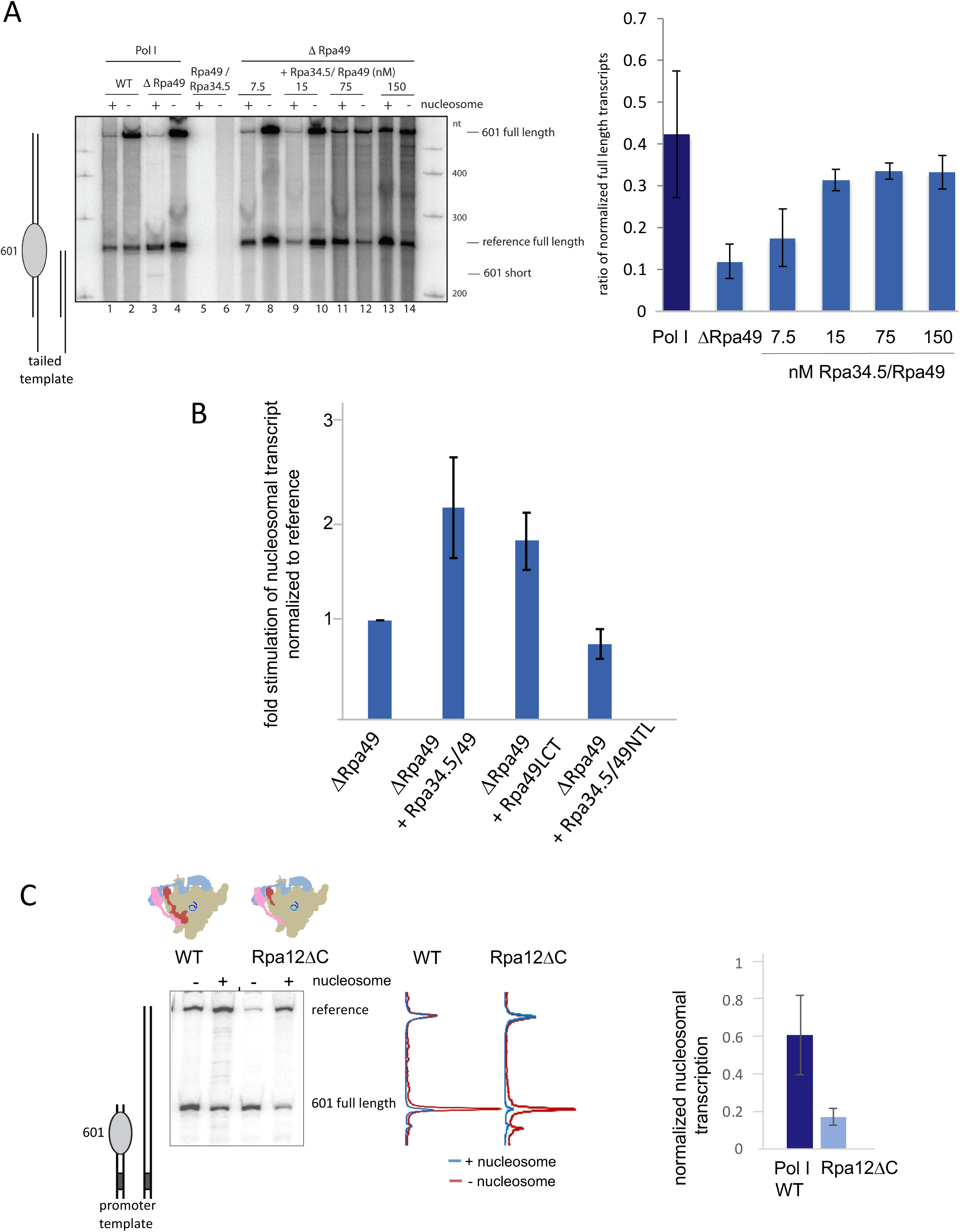
Efficient transcription through nucleosomes requires the C-terminal part of Rpa49 and the C-terminal part of Rpa12.2. A) The heterodimeric subunit Rpa34.5/Rpa49 of Pol I is required for efficient transcription through a nucleosome in tailed template assays. *Left panel* Transcription reactions were performed as in Fig. 1A using 5 nM WT Pol I (lanes 1 and 2) or 15 nM Pol I lacking subunits Rpa49 and Rpa34.5 (ΔRpa49), lanes 3 and 4). Increasing amounts of purified recombinant Rpa34.5/Rpa49 heterodimer were added to Pol I ΔRpa49 in reactions analysed in lanes 7 to 14. Reactions analysed in lanes 3 and 4 were performed using 150 nM Rpa34.5/Rpa49 heterodimer, but no polymerase. Transcripts were analyzed on a 6% polyacrylamide gel containing 6 M urea. (601 short indicates the position of a stalled transcript at the 5’end of a nucleosome; see also Appendix Fig. S2). *Right panel.* Quantification from two independent experiments. B) Stimulation of ΔRpa49 Pol I to transcribe nucleosomal templates in the presence of Rpa34.5/Rpa49 heterodimer, Rpa49LCT and Rpa34.5/Rpa49NTL. Signals of nucleosomal transcripts were normalized to the respective reference transcripts and the ratio derived from the ΔRpa49 Pol I was set to 1. The single experiments (5 respective 3 independent experiments) including experimental conditions are depicted and described in Appendix Fig. S4). For determination of the mean values the lanes showing saturated amount of transcripts were used for each set of add-in experiment. C) Efficient transcription through nucleosomes requires the C-terminal domain of Rpa12.2. *Left panel* Promoter-dependent transcription was performed for 30 min on templates with and without attached nucleosomes using 70 nM Rrn3, 20 nM CF and 4 nM Pol I WT, Pol I ΔRpa12 and Pol I Rpa12ΔC in the presence of 200 μM ATP, 200 μM UTP, 200 μM GTP, 10 μM CTP (^32^P-CTP). The cartoon on the left indicates the length of the transcripts derived from the nucleosomal and reference template. (Note, here the reference template is longer than the nucleosomal template). Transcript profiles of each lane are shown on the right. The value of the reference transcript was set to 1. *Right panel* Quantification (n=3), the ratio of nucleosomal transcripts was normalized to transcripts derived from non-nucleosomal transcription.

To find out whether the RNA cleavage-stimulating activity of Rpa12.2 contributes to transcription through nucleosomes, transcription assays were performed using WT Pol I and Pol I Rpa12ΔC. Both enzyme preparations contain Rpa34.5/Rpa49 (Appendix Fig. S3A/B). Therefore, in contrast to ΔRpa12.2 Pol I, which lacks Rpa34.5/Rpa49, Rpa12ΔC Pol I is still able to efficiently initiate transcription at the Pol I promoter together with recombinant Rrn3 and CF (Fig. 2C). Promoter-dependent transcription by Rpa12ΔC Pol I was much stronger affected by the presence of a nucleosome than transcription of WT Pol I (Fig. 2C and Appendix Fig. S2B). This suggested that - in addition to the Rpa49/Rpa34.5 heterodimer – the C-terminus of Rpa12.2 is also required for efficient passage through a nucleosome, adding a novel role to the domain. Whether this role is dependent on the cleavage activity remains to be determined.

In summary, the C-terminal domain of Rpa12.2 and the Rpa34.5/Rpa49 heterodimer both support Pol I transcription of nucleosomal templates.

### Pol I elongation rate or processivity influence transcription through nucleosomes

A physical barrier like a nucleosome may promote stalling and backtracking of the transcribing Pol and challenge resumption of efficient transcription elongation [11]. Accordingly, high processivity or elongation rate could be an important feature of a RNAP for efficient transcription of nucleosomal templates. It was previously shown that reduction of nucleotides reduces the elongation rate of RNA polymerase I and II [50,51]and that reduced elongation rate of Pol II influences nucleosomal transcription [51]. Therefore, it was tested if nucleotide reduction affects Pol I processivity and movement through *in vitro* assembled nucleosomes. When *in vitro* transcription reactions were performed in the presence of 20 μM NTPs instead of 200 μM NTPs Pol I transcription of a nucleosomal template was significantly impaired (Fig. 3A and C). Similar effects were observed, when transcription reactions in the presence of 200 μM NTPs were performed at 4°C instead of 24°C (Fig. 3B and C).

**Fig. 3.**
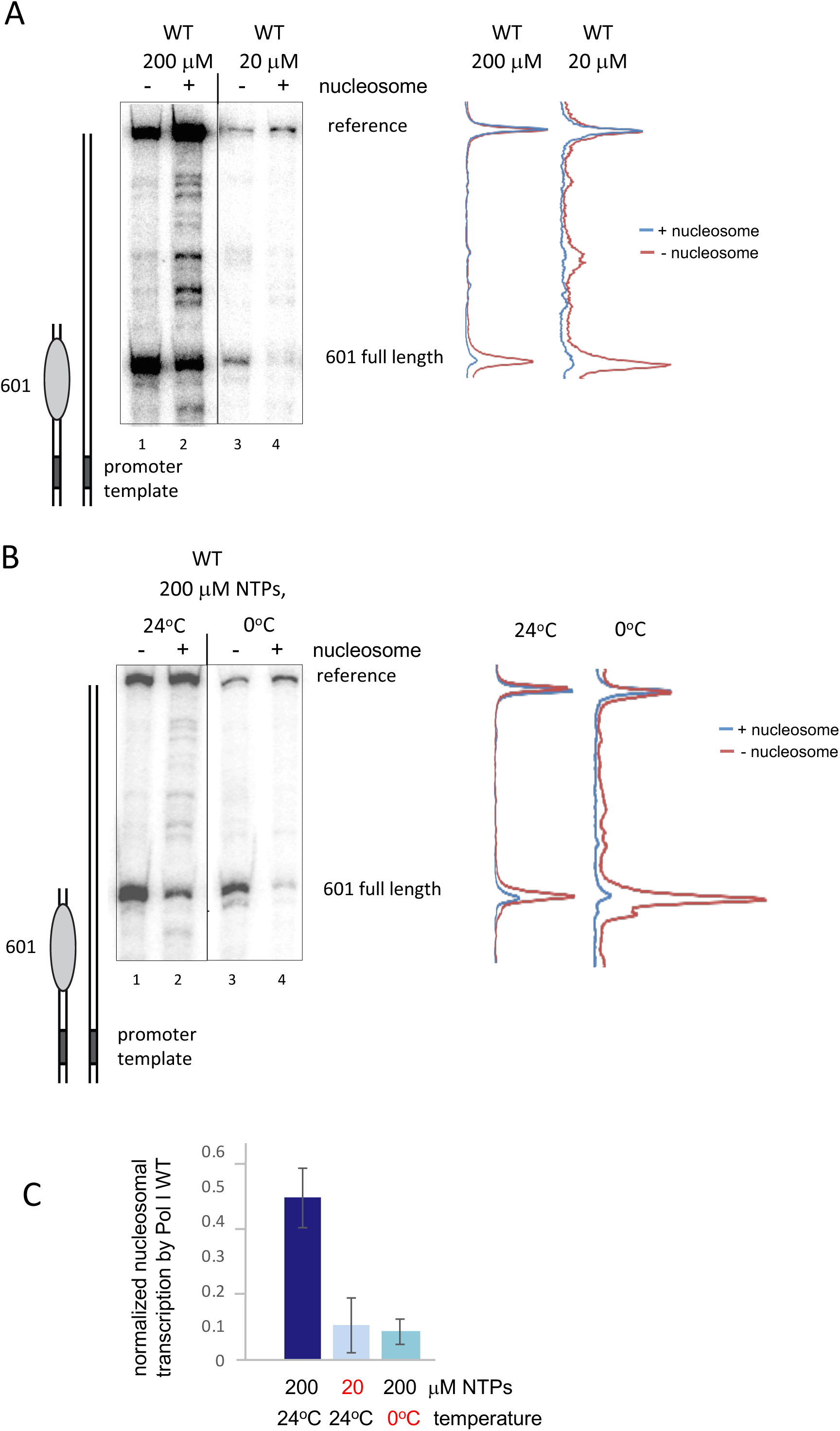
Low nucleotide concentrations and low temperature hamper efficiency of transcription through nucleosomes. A) Promoter-dependent transcription was performed on templates with and without attached nucleosomes using 70 nM Rrn3, 20 nM CF and 4 nM Pol I WT. The templates contained a 22nt long stretch without guanines after the start site on the coding strand. To achieve sufficient and comparable elongation competent Pol I complexes, transcription was started using 200 μM ATP, UTP and 10 μM α^32^P-CTP, but no GTP which should result in stalled ternary complexes 22 nt downstream of the Pol I start site. After 5 min the transcription reaction was either 10 x diluted with transcription buffer containing 200 μM ATP, UTP and 10 μM ^32^P labeled CTP and 200 μM GTP (lanes 1 and 2) or transcription buffer containing 20 μM GTP and 10 μM α^32^P-CTP but no ATP and UTP which should result in a final NTP concentration of 20 μM (lanes 3 and 4). Transcription elongation was stopped after 10 min. Transcripts were analyzed on a 6% polyacrylamide gel containing 6 M urea. B) Promoter-dependent transcription was performed on templates with and without attached nucleosomes using 70 nM Rrn3, 20 nM CF and 4 nM Pol I WT. To achieve sufficient and comparable elongation competent Pol I complexes, transcription was started using 200 μM ATP, UTP and 10 μM α^32^P-CTP, but no GTP at 24°C which should result in stalled ternary complexes 22 nt downstream of the Pol I start site. After 5 min the transcription reaction was 10 x diluted with either 24°C (lanes 1 and 2) or 4°C cold transcription buffer containing 200 μM ATP, UTP and 10 μM ^32^P labeled CTP and 200 μM GTP (lanes 3 and 4). Transcription elongation was stopped after 10 min. Transcripts were analyzed on a 6% polyacrylamide gel containing 6 M urea. C) Quantification of the experiments A and B. Signal intensities of 601 full length transcripts of Pol I WT were normalized to reference full length transcript signal intensities. Then, the ratio of the normalized 601 full length transcript signal intensities in presence and absence of the nucleosome was calculated. Pol I WT 200 μM (n=5), Pol I WT 20 μM (n=2), Pol I WT 200 μM 4°C (n=2).

To evaluate how suboptimal transcription conditions affect Pol I processivity, promoter-dependent transcription of naked DNA templates was tested at an NTP concentration of 20 μM or a reaction temperature of 4°C during chain elongation. We analysed Pol I transcript elongation using a Pol I promoter-containing template followed by 169 nt of DNA lacking guanines in the coding strand (G-less cassette, see Fig. 4A right panel). The end of the transcribed cassette of the templates was attached to magnetic beads (Fig. 4A, left panel). Transcription was initiated with Pol I, recombinant CF and Rrn3 using a transcription buffer containing 200 μM UTP, 10 μM α^32^P-CTP, and 200 μM ATP, but no GTP. Thus, Pol I could initiate transcription and transcribe the G-less cassette but was stalled when the first GTP had to be incorporated in the RNA (Fig. 4B, lanes 1 and 7). Ternary arrested complexes, composed of DNA, Pol I, and the nascent radioactively labelled RNA were washed with buffers containing either 200 μM ATP, CTP and UTP (lanes 1-6) or 20 μM ATP, CTP and UTP (lanes 7-12). This procedure removed unincorporated ^32^P-CTP and unbound proteins. Furthermore, heparin was added to inhibit (re)initiation. After addition of either 200 μM GTP (chase)(lanes 2–6) or 20 μM GTP (lanes 8-12), time course experiments were performed to follow transcript elongation (Fig. 4B). In the presence of 20 μM NTPs chain elongation was significantly slower (Fig. 4B, compare lanes 8-10 with lanes 2-4) and only up to 60% of the stalled transcripts at the G-less cassette could be fully extended (Fig. 4D). A very similar result was obtained when RNA chain elongation was performed in the presence of 200 μM NTPs, but at 4°C instead of 24°C (Fig. 4C). At lower temperature only about 60% transcripts could be fully extended (Fig. 4D) and the elongation rate was significantly lower compared to the elongation rate at room temperature (Fig. 4C, compare lanes 2-4 with lanes 7-9). Using the same experimental set-up, processivity of Pol I mutants lacking lobe binding subunits was tested in direct comparison with WT Pol I (Fig. 4E). Under optimal *in vitro* transcription conditions, ΔRpa49 Pol I or Rpa12ΔC Pol I extended only 50-60% of the stalled transcript to the full length. Interestingly, under these optimal transcription conditions, the elongation rate of the mutants was not as significantly reduced as using Pol I WT at reduced NTP levels or at lower temperature (compare time points 15 and 30 seconds, Fig. 4D and 4E). This finding suggests that rather the reduced processivity in the absence of lobe binding subunits affects movement through nucleosomes. In summary, our data suggest that the high processivity of the Pol I enzyme correlates with its ability to overcome a nucleosomal barrier.

**Fig. 4.**
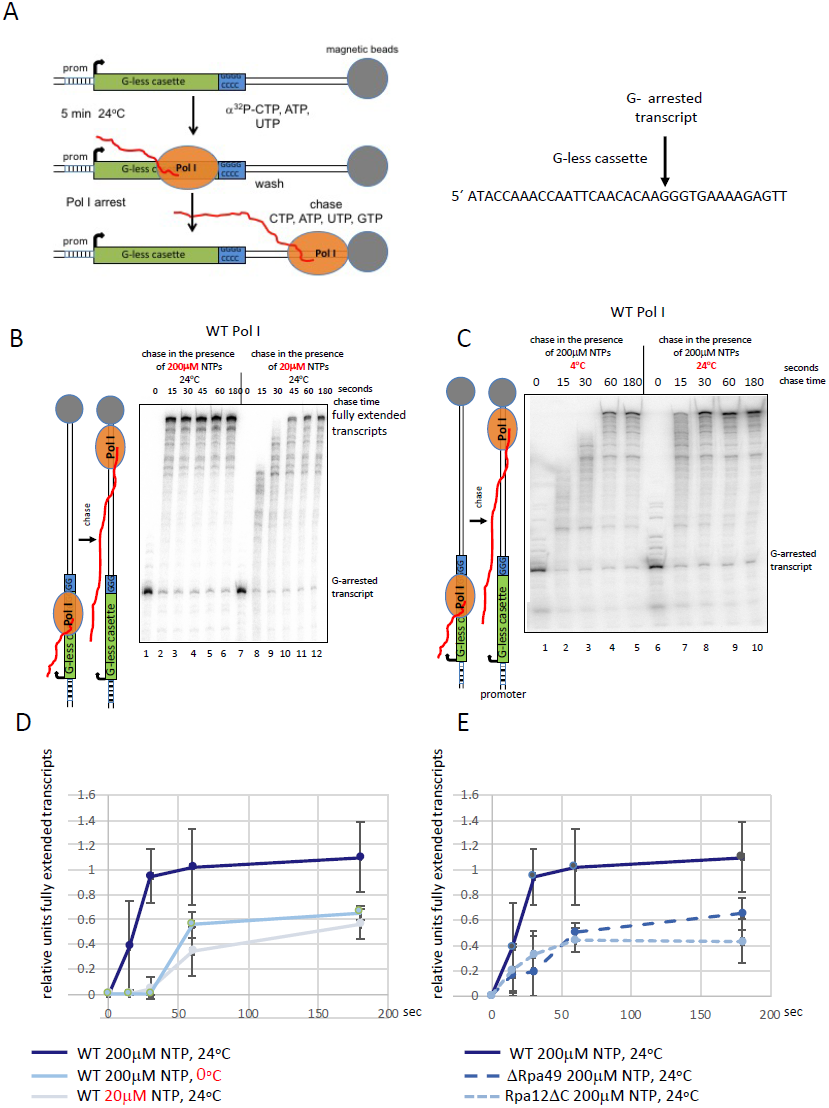
Low nucleotide concentrations, low temperature and depletion of Pol I lobe binding subunits decrease Pol I processivity. A) Experimental design. Promoter-dependent transcription was performed for 10 min on template PIP_G-extend 2kb attached to magnetic beads using 5 nM Pol I, 70 nM Rrn3 and 20 nM CF in the presence of 200 μM ATP, 200 μM UTP, 10 μM CTP (^32^P-CTP) but no GTP. Transcription was arrested after 169 nt when the first GTP should be incorporated (see Fig. 4B, lanes 1 and 7; Fig. 4 C, lanes 1 and 6). An enlarged section of the G-less cassette containing the first 3 guanosines to be incorporated is depicted on the right. The paused ternary complexes were washed with buffers containing ATP, CTP and UTP, but lacking ^32^P-CTP and GTP. Elongation kinetics were performed after addition of GTP. B) Influence of nucleotide concentration on elongation processivity. After 5 min incubation at 24°C, the arrested ternary complexes were washed twice with excess transcription buffer containing 20 μM ATP, 20 μM UTP and 20 μM CTP but no GTP. Washed ternary complexes were chased using either 200 μM NTPs (lanes 2 - 6) or 20 μM NTPs (lanes 8 – 12) at 24°C. At the time points indicated 25 μl aliquots were withdrawn and injected in 200 μl 0.6% SDS, 0.3 M NaCl, 5 mM EDTA, 0.5 mg/ml Proteinase K, which immediately stopped rRNA synthesis. Transcripts were analyzed on a 6% polyacrylamide gel containing 6 M urea and detected using a PhosphorImager. C) Influence of temperature on elongation processivity. After 5 min incubation at 24°C, the arrested ternary complexes were washed twice with ice cold excess transcription buffer containing 200 μM ATP, 200 μM UTP and 200 μM CTP but no GTP. Washed ternary complexes were chased using either 200 μM NTPs (lanes 2 - 5) at 4°C or 200 μM NTPs (lanes 67– 10) at 24°C. D) Quantification of experiments 4B an 4C. Pixel intensities of the bands of fully extended transcripts were normalized to the pixel intensities of the band of the transcript stalled at the G-less cassette at time point 0. (WT 200 μM NTP, 24°C n=4; WT 20 μM NTP, 24°C n= 4; WT 200 μM NTP, 4°C n=2) E) Pol I ΔRpa49 and Pol I Rpa12ΔC elongate stalled transcription complexes with reduced efficiency. Promoter dependent transcription reactions were performed as in 4B, (lanes 1 and 7) but using 20 nM Pol I ΔRpa49 or Pol I Rpa12ΔC. Washed ternary complexes were chased using 200 μM NTPs at 24°C. Pixel intensities of the bands of fully extended transcripts were normalized to the pixel intensities of the band of the transcript stalled at the G-less cassette at time point 0. (WT; n=4 Pol I ΔRpa49; n=3 Pol I Rpa12ΔC; n=3)

## Discussion

If elongating Pol II encounters a barrier, the enzyme stalls, backtracks and arrests [52–54]. Rapid relief of Pol II arrest requires additional factors including RNA cleavage stimulating proteins and cleavage of the 3’ end of the RNA which is displaced from the active center. Nucleosomes are a physiological relevant barrier for Pol II and the other nuclear eukaryotic RNA polymerases. In contrast to Pol II, Pol I encounters a nucleosomal template presumably only once during the cell division cycle, when nucleosomes were deposited by the replication machinery. Likely correlating with the pioneering round of transcription, nucleosomes are depleted from rRNA genes by an unknown mechanism. In summary, our data show that the heterodimer Rpa34.5/Rpa49 and subunit Rpa12.2 are required to ensure high Pol I processivity *in vitro* which correlates with efficient passage through nucleosomes. The stable association of lobe binding subunits might support rapid and processive movement through DNA-bound barriers. Whether or not the nucleosome is evicted by Pol I or the DNA is transiently uncoiled during elongation, remains to be determined. We tried to distinguish between the possibilities of eviction and re-assembly of nucleosomes, but our ensemble assays were not sensitive enough for a clear-cut result. Thus, protection of the nucleosomal template against restriction enzyme digestion was found to be very similar before and after transcription (Appendix Fig. S5). Only less than 10% template of the nucleosomal templates were transcribed in our assay. Restriction enzyme accessibility assays are in our hands not sensitive enough to reliably detect nucleosome disassembly in such a small subpopulation. Future single molecule analysis might help to answer this question.

For Pol II transcription it was described that nucleosomal DNA can recoil on the octamer, locking the enzyme in the arrested state [55]. Rapid release from the arrest requires transcription factor TFIIS and cleavage of the 3’ end of the RNA which is displaced from the active center of Pol II. TFIIS-mediated *in vitro* transcription through nucleosomes might be supported by TFIIF [18]. None of these factors remained associated with the affinity purified Pol II used in the above experiments. This explains why Pol II transcription of the nucleosomal template was almost fully abolished (Fig. 1A). In contrast, Pol I and Pol III, whose homologous subunits Rpa34.5/Rpa49, Rpa12.2 and Rpc37/Rpc53, Rpc11 resemble TFIIF and TFIIS, efficiently transcribed the same nucleosomal template. For Pol I, the C-terminal part of Rpa12.2 contributes to the ability to transcribe through nucleosomes, suggesting that the RNA cleavage activity of Pol I supports passage through nucleosomes. Whereas, deletion of the C-terminal Rpa12.2 domain showed no significant growth phenotype under normal growth conditions, deletion of the N-terminus led to growth phenotypes indistinguishable from that of a *Δrpa12* strain [34]. The lack of a growth phenotype of a *rpa12ΔC* strain was surprising, since this subdomain it is important for RNA cleavage activity and recovery from long backtracks of up to 20 nucleotides [14,34,53]. It is possible that stalled Rpa12ΔC Pol I molecules are destabilized and targeted to proteasomal degradation *in vivo*. Accordingly, there was recent evidence for proteasome-mediated degradation of Pol I stalled by the drug BMH21 [56].

In the absence of the dimerization module Rpa34.5/Rpa49NT, addition of Rpa49LCT alone improved Pol I movement through nucleosomes. Several features of Rpa49LCT may explain its stimulatory effect of Pol I processivity, which allows efficient transcription of nucleosomal templates. Rpa49LCT binds DNA, supports promoter-dependent transcription initiation and extension of RNA from an *in vitro* reconstituted transcription bubble [16,41]. Recently, structural analysis of Rpa49CT provided further insights [16,29,41,57–60]. These results, in combination with genetic data, indicated that the Rpa49 linker together with the Rpa49 tWH domain might stabilize the closed conformation of DNA-bound Pol I, thereby maintaining a narrow cleft [61,62]. It is possible that an enzyme containing Rpa49LCT, but no Rpa34.5/Rpa49NT may not enter a locked state with a 3’ RNA displaced from the active center, and, thus, supports high processivity.

For passage through nucleosomes uncoiling of the template from the histone octamer is required to maintain the polymerase in the transcriptionally competent state [52]. Physical and chemical parameters of *in vitro* transcription reactions, like nucleotide concentration, which influence transcription rate, might alter histone transfer and the fate of transcribed nucleosomes [51]. Accordingly, nucleotide concentration and/or transcription rate can change the interaction between an elongating Pol II and an assembled nucleosome. Our results suggest that both nucleotide levels and elongation rate affect Pol I interaction with nucleosomes and lead to a less processive enzyme in the presence of a nucleosome. Reduced Pol I processivity could have several reasons. It is possible that *i)* Pol I dissociation from the DNA template is facilitated, *ii)* Pol I is more often stalled because of R-loop formation [63,64] or *iii)* Pol I is backtracked at frequent intrinsic pausing from which the enzyme cannot resume elongation. However, the underlying reason for the loss of processivity of the different Pol I mutants remains elusive at this time.

RNAP movement through nucleosomes can also be supported by factors which weaken the interaction between nucleosomes and DNA in context with transcription. Histone chaperone FACT or elongation factors Spt4/5 or Paf1c are examples of Pol II factors that facilitate DNA unwrapping from H2A/H2B dimers to partially disassemble nucleosomes [48,49,65,66]. The same factors were also reported either to support Pol I elongation [67–69] or passage through nucleosomes [13]. None of these factors was present in the assays performed in this study. However, it is possible that *in vivo* Pol I subunits and elongation factors cooperate to support transcription through nucleosomes.

The finding that mutant strains expressing Pol I enzymes lacking either Rpa34.5/Rpa49 or RPA12.2 are viable support this point of view [27,33,70]. Obviously, mutant polymerases can still transcribe nucleosomal templates *in vivo* which suggests that other factors of the Pol I transcription machinery contribute to this process. Such candidates could be the above mentioned FACT [13] or Paf1c [68] which both interact with chromatin.

Our studies that the lobe binding subunits Rpa34.5/Rpa49 and Rpa12.2 facilitate passage through nucleosomes point to a possible cooperation between these subunits. Recent structural investigations suggest that the C-terminal part of Rpa12.2 (Rpa12.2CT) may adapt different positions within the Pol I enzyme. Accordingly, a dynamic position of Rpa12.2CT in the funnel of subunit Rpa190 depends on the transcriptional status of the elongating Pol I and might be a prerequisite to overcome transcriptional barriers [19,20,26,29,29,41,57,60,71,72]. Furthermore, it was suggested that competition between Rpa12.2CT and the heterodimer Rpa34.5/Rpa49 for binding of the Pol I lobe might result in the reversible binding of the heterodimer which could contribute to modulate Pol I activity [62]. Such a dynamic interaction could also influence movement through nucleosomes. However, this hypothesis would disagree with ChIP experiments indicating that Rpa34.5/Rpa49 remain associated with elongating Pol I [28].

Hence, the next challenges are to understand whether and how the lobe binding subunits cooperate in detail and to elucidate the mechanisms how they support stalled Pol I complexes to bypass transcription obstacles and resume elongation. This will contribute to our knowledge how the specific subunit composition of Pol I leads to its functional specialisation, namely to transcribe the rRNA genes as efficiently as possible.

## Acknowledgements

We are grateful to members of the chair Biochemistry III for critical discussion and thank Dr. Jorge Perez-Fernandez for providing us with yeast strains.

